# Histone H3K27 methylation perturbs transcriptional robustness and underpins dispensability of highly conserved genes in fungi

**DOI:** 10.1101/2021.05.21.445187

**Authors:** Sabina Moser Tralamazza, Leen Nanchira Abraham, Benedito Corrêa, Daniel Croll

## Abstract

Epigenetic modifications are key regulators of gene expression and underpin genome integrity. Yet, how epigenetic changes affect the evolution and transcriptional robustness of genes remains largely unknown. Here, we show how the repressive histone mark H3K27me3 influences the trajectory of highly conserved genes in fungi. We first performed transcriptomic profiling on closely related species of the plant pathogen *Fusarium graminearum* species complex. We determined transcriptional responsiveness of genes across environmental conditions to determine expression robustness. To infer evolutionary conservation of coding sequences, we used a comparative genomics framework of 23 species across the *Fusarium* genus. We integrated histone methylation data from three *Fusarium* species across the phylogenetic breadth of the genus. Gene expression variation is negatively correlated with gene conservation confirming that highly conserved genes show higher expression robustness. Furthermore, we show that highly conserved genes marked by H3K27me3 deviate from the typical housekeeping gene archetype. Compared to the genomic background, H3K27me3 marked genes encode smaller proteins, exhibit lower GC content, weaker codon usage bias, higher levels of hydrophobicity and are enriched for functions related to regulation and membrane transport. The evolutionary age of conserved genes with H3K27me3 histone marks falls typically within the origins of the *Fusarium* genus. We show that highly conserved genes marked by H3K27me3 are more likely to be dispensable for survival. Lastly, we show that conserved genes exposed to repressive H3K27me3 marks across distantly related fungi predict transcriptional perturbation at the microevolutionary scale in *Fusarium* fungi. In conclusion, we establish how repressive histone marks determine the evolutionary fate of highly conserved genes across evolutionary timescales.

## Introduction

Highly conserved genes are ensuring housekeeping functions essential for the survival of the organism (She et al. 2009). Beyond sequence conservation, highly conserved genes are often constitutively expressed. Changes in gene expression is a major factor associated with the evolutionary trajectory of genes. In yeasts, gene expression levels are negatively correlated with the evolutionary rate of gene sequences (Pál et al. 2001; Drummond et al. 2006). The correlation, also termed E-R correlation, was later confirmed across the tree of life including for bacteria (Rocha and Danchin 2004; Drummond et al. 2006) Metazoa (Krylov et al. 2003) and plants (Ingvarsson 2006). Both empirical and theoretical approaches support that the E-R correlation stems from the fact that highly expressed genes show stronger functional constraints. These include purifying selection acting against deleterious mutations affecting translation, protein folding and maladaptive protein interactions (Zhang and Yang 2015). Despite the broad evidence that gene expression levels are strongly associated with the rate of protein evolution across species, how the association arises through genetic and epigenetic modifications over shorter evolutionary time spans remains largely unknown.

Gene expression in eukaryotes is governed largely by higher-order chromatin structure (Woodcock and Ghosh 2010). Chromatin is determined by the nucleosome, which contains a histone octamer attached to a stretch of nuclear DNA (Luger et al. 1997). Histones and DNA are constantly being modified by various proteins catalyzing enzymatic activities, such as phosphorylation, acetylation, methylation, ubiquitination, and O-GlcNAcylation. The joint effect of these post-translational modifications regulates and stabilize gene expression during the cell cycle and development while facilitating responses to environmental stimuli (Youn 2017). For example, cytosine methylation is a covalent DNA modification associated with transcriptional responses to biotic stress (Dowen et al. 2012) and plays a key role in genomic defenses against selfish genetic elements in plants (Cui et al. 2013), mammals (Smith and Meissner 2013) and fungi (Zhou et al. 2001). Histone modifications also dynamically remodel chromatin structure to activate or repress gene expression following intrinsic and extrinsic signals (Flavahan et al. 2017). Specific histone modifications such as the methylation of H3K9 residues are frequently associated with transcriptionally silent heterochromatin, while acetylation and methylation of H3K4 and H3K36 residues are hallmarks of transcriptionally active euchromatin (Gates et al. 2017). The H3K27me3 modification plays a particularly important role by underpinning facultative heterochromatin allowing for responsive gene expression regulation (Trojer and Reinberg 2007).

The H3K27 residue is methylated by the protein Polycomb repressive complex 2 (PRC2) (Qian et al. 2006). The canonical formation of the complex constitutes three core domains KMT6/EZH2, EED, SUZ12 which are conserved in multiple taxa such as vertebrates, insects, plants, and fungi (Ridenour et al. 2020). H3K27 marks are often associated with transcriptional repression in gene-rich chromosomal regions (Jamieson et al. 2013), control of developmental stages in higher organisms such as plants and flies (Schwartz and Pirrotta 2007) and genome instability in fungi (Janevska et al. 2018; Möller et al. 2019). Due its importance for both gene regulation and genome integrity, H3K27me3 is one of the best-studied post-translational marks in fungi. H3K27me3 marks show a wide range of abundance from ~7% in the *Neurospora crassa* genome to ~30% in *Fusarium graminearum* (Freitag 2017). In the budding yeast *Saccharomyces cerevisiae* and fission yeast *Schizosaccharomyces pombe*, H3K27 methylation is absent due to the loss of the Polycomb complex (Connolly et al. 2013). Experimentally induced loss of H3K27 methylation has highly variable impacts. Inactivation of the Polycomb complex (*i.e.* PRC2) in *Fusarium* fungi causes growth and developmental defects, as well as sterility (Freitag 2017). However, the inactivation has no overt effect in *N. crassa*. Yet deletion of the methyltransferase *set-1* underlying H3K4me3 causes promiscuous spread of H3K27me3 across the genome and severe fitness costs (Jamieson et al. 2013; Kronholm et al. 2016).

Across fungi, H3K27me3 regulates the expression of rapidly evolving genes encoding virulence factors and biosynthetic pathways of specialized metabolites (Connolly et al. 2013; Jamieson et al. 2013). How facultative heterochromatin such as governed by H3K27 methylation impacts the evolution and transcriptional robustness of more conserved genes remains largely unknown. Studies focused on deep evolutionary timescales (Fair et al. 2020) and a small number of highly conserved orthologues across taxa. However, identifying causal factors underlying among species transcriptional variability is challenging because of environmental heterogeneity and niche differentiation (Dötsch et al. 2015). Furthermore, technical noise arising from different experimental setups renders comparisons difficult (De Jong et al. 2019) and uncertainty about orthology inference and functional divergence add additional uncertainty. In contrast, closely related fungi provide ideal models to analyze how repressive histone marks underpin the robustness of expression and sequence conservation across species. The ability to access transcriptional variability in identical experimental settings in species sharing colinear genomes, similar habitats and life cycles is a key advantage to make robust inferences about causal factors.

Fungi of the genus *Fusarium* are important pathogens of crops causing a wide range of diseases (Summerell 2019). Originally described as a single species (O’Donnell et al. 2004), the *Fusarium graminearum* species complex (FGSC) comprises 16 recognized species (O’Donnell et al. 2008). All species can cause Fusarium head blight in cereals and hence have strongly overlapping host ranges (van der Lee et al. 2015). FGSC are all closely related (Walkowiak et al. 2016; Tralamazza et al. 2019) but encode a vast and variable repertoire of specialized metabolites (Tralamazza et al. 2019). The genome of *F. graminearum* has one of the most strongly affected gene bodies in terms of repressive H3K27me3 marks among fungi (46% of genes) (Freitag 2017) making FGSC an ideal model to analyze epigenetic control of gene expression.

To investigate the impact of repressive histone marks on gene expression robustness and protein conservation across species, we performed genome-wide expression analyses of five FGSC members including the reference genome strain of *F. graminearum* (PH-1) (Cuomo et al. 2007). We analyzed transcriptional responsiveness across two environmental conditions including the infection of the wheat host and a nutrient-rich growth medium. To infer evolutionary conservation of coding sequences, we used a comparative genomics framework of 23 species across the genus of *Fusarium* spanning approximately 110-420 million years of evolution (Summerell et al. 2010). We integrated histone methylation data from three *Fusarium* species across the phylogenetic breadth of the genus. We found that gene expression variation is negatively correlated with gene conservation across the genus confirming that highly conserved genes show higher gene expression robustness. Genes silenced through H3K27me3 histone marks in any of the species showed lower expression levels and higher gene expression variation among species. Marked genes encoded for on average smaller proteins, showed lower GC content, codon usage bias and were enriched for functions related to regulation and membrane transport. Estimates of the evolutionary age of conserved genes showed that genes with H3K27me3 histone marks are of much more recent origin than unaffected genes. Lastly, we show that highly conserved genes marked by H3K27me3 are more likely to be dispensable.

## Material and Methods

### Infection Assay

RNA-seq analyses were performed on fungal mycelium grown in culture and *in planta* for five species of the FGSC (*F. graminearum, F. meridionale, F. cortaderiae, F. asiaticum* and *F. austroamericanum*) (Figure 1A). Each species was grown in a Petri dish containing V8 agar medium for four days at 25°C. Following the culturing, the mycelium was transferred to a 100 ml mung bean liquid medium and agitated at 170 rpm for five days at 25°C (Zhang et al. 2012). Next, the medium was filtered, and spores counted using a haemocytometer. A 10^6^ per ml spore solution was prepared for the following infection procedure. For *in vitro* assays, cultures were inoculated from spore solutions in yeast sucrose agar medium for 72h at 25°C until RNA extraction. For *in planta* assays, wheat coleoptiles with intermediate susceptibility to Fusarium head blight (cultivar CH Combin, harvest 2018-2019) were infected with each species individually according to (Zhang et al. 2012). Briefly, wheat seeds were soaked in sterilized water for germination in a culture chamber at 25°C with a 12h white light cycle and 93% humidity. After 3 days of germination, the tip of the coleoptile was cut and 10μl of the spore solution was used for inoculation. Coleoptiles were collected 72h after inoculation (approximately 0.4 mm lesion size) and processed for RNA extraction. All *in planta* assays were performed in triplicates and each replicate was composed of a pool of 25 infected seedlings to obtain sufficient material and homogenize infection conditions.

**Figure 1.**
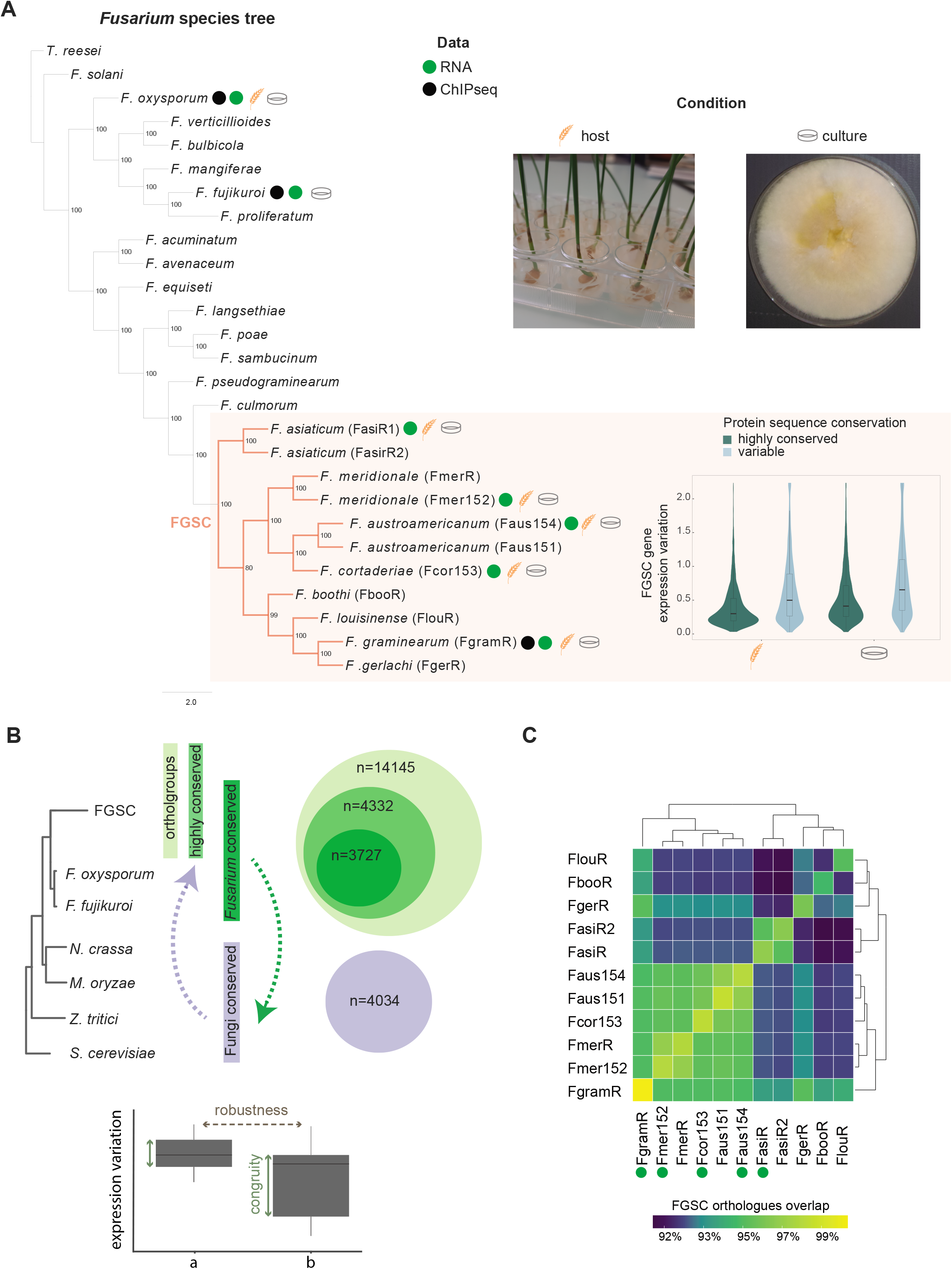
The *Fusarium graminearum* species complex (FGSC). A) Phylogenomic tree of *Fusarium* species was inferred from a coalescence-based analysis of 4192 single-copy orthologues (Tralamazza et al. 2019). Values indicate support from 100 bootstrap replicates. *Trichoderma reseei* was used as an outgroup. Colored circles indicate data type utilized in the current study. Host and culture medium describe the two environmental conditions. B) Outline of the datasets used in the study. Colored dashed arrows refer to the evolutionary fate inquiry across taxon. Boxplot refers to the expression robustness and expression congruity defined in the study. Green solid arrow indicates comparison within the same group (*e.g* group a), grey dotted arrow indicate comparison between groups (*e.g*. a vs. b). C) Percentage of orthologues shared among FGSC members. The green circle indicates RNA-seq analysis.

### RNA extraction and sequencing

RNA extraction was performed using the NucleoSpin RNA Plant and Fungi kit (Macherey□Nagel GmbH & Co. KG, Düren, Germany) according to the manufacture’s recommendation. The RNA quality was assessed using a Qubit (Thermo Fisher Scientific, Waltham, USA) and an Agilent Bioanalyzer 2100 (Agilent Technologies, CA, USA). The NEB Next® Ultra™ RNA Library Prep (NEB, CA, USA) kit based on the polyA method was used for RNA library preparation. Samples were sequenced on a NovaSeq 6000 system (Illumina Inc.), CA, USA and 150 bp paired-end reads were generated. Library preparation and sequencing was performed by Novogene Co., Ltd, Beijing, China.

### RNA-seq read alignment and transcript quantification

Illumina raw reads were trimmed and filtered for adapter contamination using Trimmomatic v. 0.32 (parameters: ILLUMINACLIP:Trueseq3_PE.fa:2:30:10 LEADING:3 TRAILING:3 SLIDINGWINDOW:4:15 MINLEN:36) (Bolger et al. 2014). Filtered reads were aligned using Hisat2 v. 2.0.4 with default parameters (Kim et al. 2019) to the *Fusarium graminearum* PH-1 reference genome (King et al. 2017). Mapped transcripts were quantified using HTSeq-count (Anders et al. 2015). Read counts were normalized based on the trimmed mean of M-values (TMM) method using the calcNormFactors option. To account for gene length, we calculated reads per kilobase per million mapped reads (RPKM) values using the R package edgeR (Robinson et al. 2010). Gene expression robustness (*i.e.* gene expression variation) among the five species of the FGSC was calculated using the coefficient of variation (CV) based on the ratio of the standard deviation to the mean (Figure 1B). To compare gene expression levels among the more distant species, we normalized and ranked gene expression values using the *percent_rank* function of the *dplyr* R package (Wickham et al. 2019). To compare expression congruity (*i.e.* expression range within a group), we calculated the interquartile range distribution, which reports the difference between the 25th and 75th percentiles (Figure 1B).

### Orthology and sequence conservation analyses

Gene orthology within the *Fusarium* genus was predicted using Orthofinder v2.4.0 with default parameters (Emms and Kelly 2019). The analysis included eight distinct species (for 11 genomes in total) for the FGSC and 15 additional species of the genus. Protein conservation was assessed as the mean of the pair-wise amino acid sequence identity based on the top BLASTP (Camacho et al. 2009) hit between protein sequences of *F. graminearum* (PH-1) against the 25 other *Fusarium* genomes. We classified the underlying gene as highly conserved if an ortholog of *F. graminearum* reference genome was conserved in at least 25 of the 26 genomes and the mean of the amino acid identity was >70% among orthologs. Only single copy genes in the FGSC were retained. Conversely, orthologs failing the above criteria were classified as variable. To infer orthology and duplication events among more distantly related fungi, we analyzed the following set of genomes using Orthofinder with default parameters: *F. graminearum, F. oxysporum, Neurospora crassa, Magnaporthe oryzae, Zymoseptoria tritici* and *Saccharomyces cerevisiae*. The outline of the datasets used in the study based on orthology conservation is available in Figure 1B. To infer the age of highly conserved genes in *Fusarium*, we used the *F. oxysporum* gene set available from the Geneorigin database covering 565 species from Ensembl and Ensembl Genomes databases following the protein family-based pipeline based on the Wagner parsimony algorithm (Tong et al. 2020).

### Chromatin immunoprecipitation (ChlP-seq)

*F. graminearum* PH-1 strain ChIP-seq and RNA-seq datasets of the histone modifications H3K27me3, H3K36me3, H3K4m3 and H3K4m2 from fungal mycelium grown in culture were retrieved from the NCBI SRA database (accession PRJNA221153) (Connolly et al. 2013). Chip-seq raw reads were trimmed with Trimmomatic v. 032 (Bolger et al. 2014) (parameters: ILLUMINACLIP:Trueseq3_SE.fa:2:30:10 LEADING:3 TRAILING:3 SLIDINGWINDOW:4:15 MINLEN:20) and mapped to the *F. graminearum* reference genome using Bowtie2 v. 2.4.0 (Langmead and Salzberg 2012). Alignment bam files were converted using BEDtools v.2.30.0 (Quinlan 2014) and peak calling was performed using the makeTagDirectory and findPeak (parameters: -style histone -region -size 1000 -minDist 1000 -C 0) programs included in Homer v.4.11 (Heinz et al. 2010). Peak calls from replicates were merged with BEDtools intersect. Peak coverage was calculated with BEDtools coverage. Similarly, gene coverage was analyzed with BEDtools intersect based on the *F. graminearum* reference genome annotation. For RNA-seq analyses of *F. graminearum* (Connoly, et al. 2013) and *F. oxysporum* (Fokkens et al. 2018), raw reads were analyzed as described above. Similarly, ChIP-seq datasets for different species were analyzed following the same protocol as described above using the matching reference genome (Supplementary Table 1). For *F. fujikuroi*, we used gene expression data based on a *F. fujikuroi* specific NimbleGen microarray analysis provided by the authors (Studt et al. 2016). Further information on the datasets used are available in Supplementary Table 1.

### Functional enrichment analyses and codon usage

Gene ontology (GO) term enrichment analyses were performed using the Fisher’s exact test based on gene counts with the *topGO* R package (Alexa and Rahnenfuhrer, 2019) and plotted using the *GOplot* R package (Walter et al. 2015). Codon adaptation usage (CAI) between marked and unmarked and genes were calculated using the CAI software of EMBOSS (http://www.ch.embnet.org/EMBOSS/) package. We used the codon usage table for *F. graminearum* (PH-1) available from the Kazusa DNA Research Institute (Nakamura et al. 2000). Hydrophobicity (GRAVY) and aromatic scores were calculated using the CodonW software v. 1.4.4 (Peden, 2000). The GRAVY score represents the average hydrophobic index across all amino acids of a predicted protein. The aromatic score was calculated as the proportion of aromatic amino acids.

### Repeat-induced point mutations and data analyses

To detect signatures of repeat-induced point mutations (RIP), we used the software The RIPper (Van Wyk et al. 2019) on the *F. graminearum* (PH-1) genome with a window size of 1000 bp and a 500 bp step size. Kendall tau-β rank correlation tests were visualized using the *corrplot* R package (Wei et al. 2017). Genome-wide expression heatmaps were generated using the *pheatmap* R package (Kolde 2017). Upset diagrams were created with the *UpSetR* R package (Conway et al. 2019). Other figures were produced using the *ggplot2* R package (Wickham 2011).

### Data availability

All access information for data generated in this study or retrieved from public databases is described in Supplementary Table 1. RNA-Seq raw reads have been deposited at NCBI SRA database under the accession number PRJNA542165.

## Results

### Robustness of transcriptomic responses among closely related species

We performed transcriptomic profiling on closely related fungi belonging to the *Fusarium graminearum* species complex (FGSC). The group of fungi include crop pathogens infecting mainly wheat. The highly overlapping habitat and lifestyle together with the recent history of speciation make FGSC highly suitable to assess the impact of repressive histone modifications on gene conservation and expression robustness. We first assessed the degree of protein sequence conservation within the complex using orthology and find that 82.5% of genes are shared among all the members analyzed (Figure 1C). Then, we expanded the orthology analysis to 23 species covering the entire genus *Fusarium* and find 4332 highly conserved genes (orthologs of *F. graminearum* conserved in at least 25/26 *Fusarium* genomes and with > 70% mean amino acid identity) and 9813 variable genes (genes which do not meet the former criteria) across the *Fusarium* species. To assess gene expression robustness under standardized environmental conditions, we generated RNA-seq data for five FGSC members during infection of wheat and growth on nutrient-rich medium (Figure 1A). The transcriptomic data showed high reproducibility (Supplementary Figure 1). We detected reliable expression in ~78% of all genes (*n =* 11,136 out of 14,145). Most genes (90.7%) were transcribed in all five species during wheat infection and on growth medium (84.8%; Supplementary Table S2). Highly conserved genes showed higher gene expression robustness compared to variable genes (Figure 1A).

### Chromatin landscape of the *F. graminearum* genome and expression robustness

Following the association of gene conservation and transcriptional robustness within FGSC, we analyzed links between chromatin modification marks and transcription profiles of individual genes (Figure 2). For this, we assessed the occupancy of histone modifications in gene body regions of the reference genome *F. graminearum* (Connolly et al. 2013). As expected, euchromatin marks (*i.e.* H3k4me2/me3) are dominating gene body regions. Facultative heterochromatin marks (*i.e.* H3K27me3) are enriched in variable genes compared to highly conserved genes (Supplementary Figure 2A). Interestingly, the distribution of gene body H3K27me3 marks is strongly bimodal (Supplementary figure 2B). Hence, we categorized genes having either >50% covered by H3K27me3 marks or less. Transcriptional profiling of genes within the species complex revealed strong genome-wide associations between gene expression levels, expression robustness, protein sequence conservation and H3K27me3 gene marks (Figure 2A). As expected, we found that highly conserved unmarked genes were the most highly expressed genes. Remarkably, we found that highly conserved marked genes were less repressed (median 11.42 RPKM, with 11.6% showing no transcription) than less conserved marked genes (median of 0.83 RPKM, with 42.3% showing no transcription; Figure 2B). Furthermore, we found that genes marked by H3K27me3 showed much lower gene expression robustness (expression variation) compared to unmarked genes independent of protein sequence conservation (Figure 2B).

**Figure 2.**
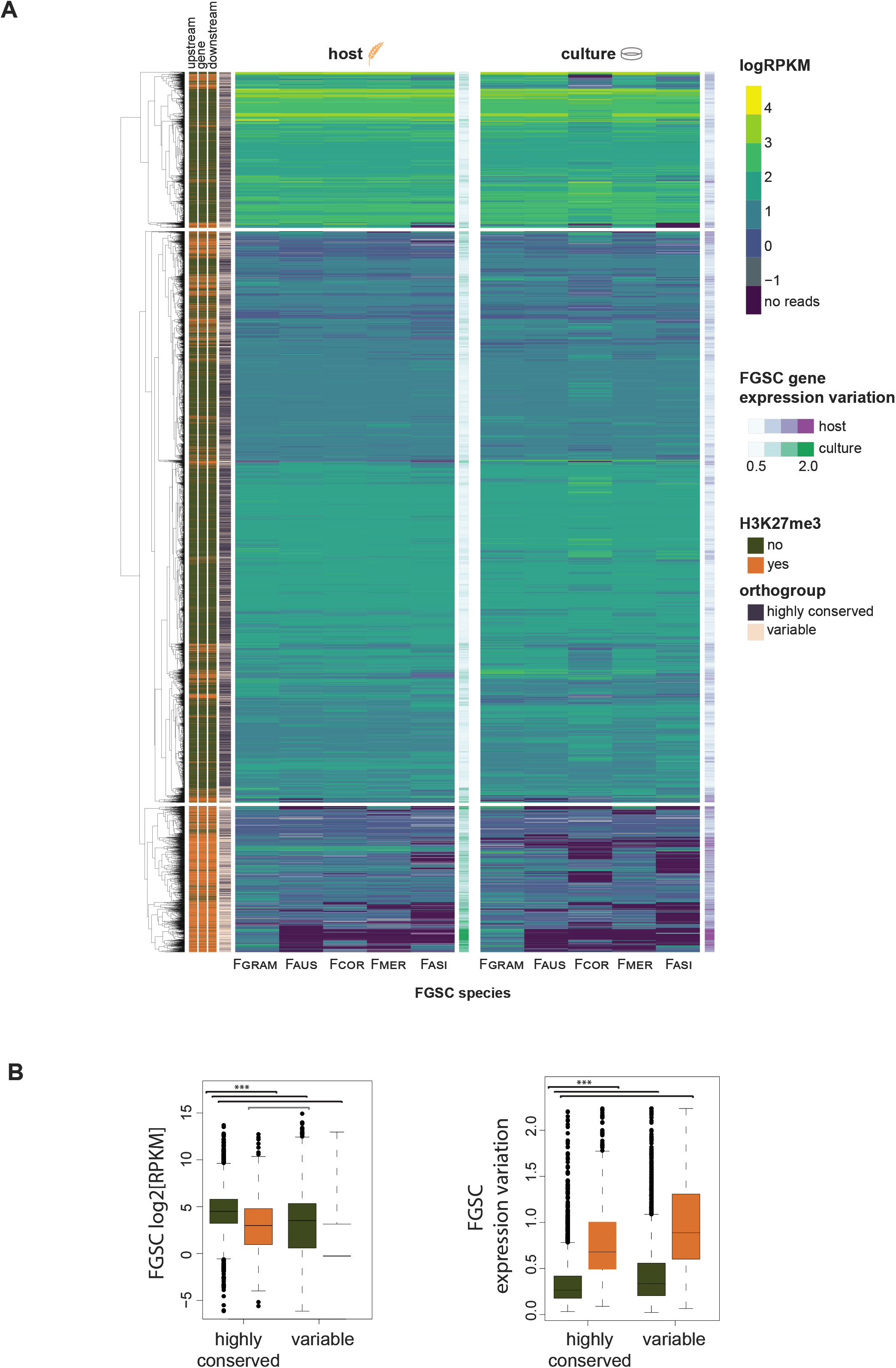
Genome-wide expression and expression variation profile of the *Fusarium graminearum* species complex (FGSC). A) Transcriptome analysis of the FGSC based on hierarchical clustering. Host and culture medium describe the two environmental conditions. Green and orange colors indicate the presence or absence of H3K27me3 in the gene body, as well as 1 kb upstream and downstream of genes. B) Mean expression and gene expression variation of highly conserved and variable genes during host infection in the FGSC. Fgram-*F. graminearum*, Faus – *F. austroamericanum*, Fcor – *F. cortaderiae*, Fmer-*F. meridionale*, Fasi-*F. asiaticum*.

To identify general patterns how H3K27me3 marks impact sequence conservation and gene expression robustness, we expanded the analyses to 4332 highly conserved genes of the *Fusarium* genus. Protein sequences of *F. graminearum* share a 98.8% mean amino acid identity with orthologs in the genus. We find support for a general E-R correlation (Figure 3A). Protein sequence conservation shows a significant positive correlation (*p*-value < 0.0001) with gene expression levels within species (*F. graminearum* on host; Kendall tau = 0.27), or between species (among FGSC members on host; Kendall tau = 0.29). We found that gene expression variation was negatively correlated (*p*-value < 0.0001) with protein sequence conservation (Kendall tau = −0.24 and −0.19 of FGSC members on host and in culture, respectively; Figure 3A). We found a strong positive correlation with histone H3K27me3 marks and gene expression variation (*i.e.* lower expression robustness; Kendall tau = 0.36 during infection and culture condition; *p*-value < 0.0001; Figure 3A).

**Figure 3.**
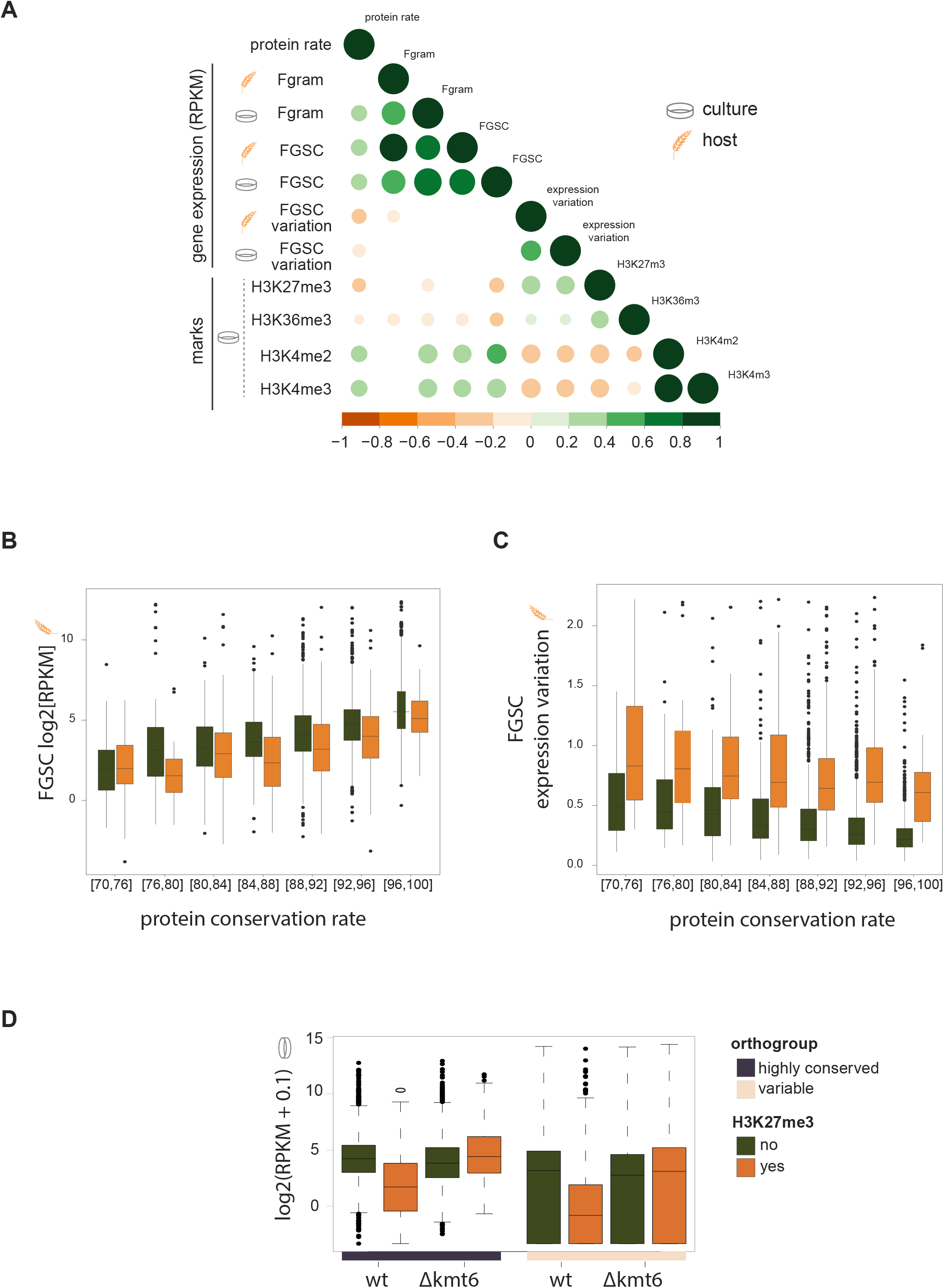
Analyses of gene expression and gene body histone methylation marks among *Fusarium graminearum* species complex (FGSC) A) Correlation plot between marks and conditions. Test was performed between paired samples, using Kendall’s *tau* method to estimate a rank-based measure of association. Circles indicate *p*-values < 0.0001. Circle size and color indicate degrees of correlation. Culture and host describe growth conditions. Fgram – *F. graminearum*. FGSC – mean value of the *F. graminearum* species complex. FGSC variation – coefficient of variation based on each species-specific RPKM values. B) Mean gene expression of highly conserved genes in the FGSC during host infection based on protein sequence conservation. C) Mean expression variation of highly conserved genes during host infection based on protein sequence conservation. Orange and green colors indicate the presence and absence of H3K27me3 in the gene body, respectively. D) Gene expression analysis of the *F. graminearum* wild type strain (wt) and mutant lacking the methyltransferase gene kmt6 (Δkmt6) assessed under culture conditions (Connolly et al. 2013). Grey and light pink refer to the degree of gene conservation.

Given the likely impact of H3K27me3 marks on expression robustness, we focused on gene expression variation among members of the FGSC. Higher gene expression is strongly associated with higher protein sequence conservation both on the host and under culture conditions (Figure 3B; Supplementary Figure 3A). Highly conserved genes (*i.e.* shared among 25/26 species and >70% identity) with H3K27me3 marks (*n* = 673) show broad transcriptional repression (Figure 3B). However, highly conserved genes show a similar range of expression (*i.e.* congruity) between different conservation rates as unmarked genes (*n* = 3659; Supplementary Figure 3B). Highly conserved genes marked by H3k27me3 are on average more divergent than unmarked genes (90.2% and 93.6% mean protein identity, respectively: Supplementary Figure 3D). Next, we analyzed gene expression robustness among members of the FGSC. Higher protein conservation was associated with reduced gene expression variation (*i.e.* higher robustness) both on the host and in culture condition (Figure 3C; Supplementary Figure 3B). We also found a positive association between protein conservation and expression congruity (*i.e.* expression range within the same category; Supplementary Figure 3C). Which supports the idea that highly conserved genes are under selection to retain transcriptional activity largely independent of the environment (*i.e.* robustness). Next, we examined the impact of H3K27me3 on robustness and found that marked genes showed significantly higher levels of gene expression variation (Figure 3C) and reduced constraints on gene expression congruity (Supplementary Figure 3B). Marked genes also show no significant association with protein conservation (Kendall tau = −0.05; *p*-value = 0.05) in contrast to unmarked genes (Kendall tau = −0.20; *p*-value < 2.2e-16).

To investigate if the loss of expression is a direct consequence of H3K27me3 marks, we analyzed RNA-seq and ChIP-seq dataset of a *F. graminearum kmt6* mutant strain (Connolly et al. 2013). The mutant lacks the methyltransferase enzyme KMT6 responsible for H3K27me3 marks in fungi (Freitag 2017). We found that highly conserved genes marked by H3K27me3 in the wild type background had expression levels similar to unmarked H3K27me3 genes in the *kmt6* mutant background. The highly conserved genes marked by H3K27me3 in the wild type background also showed higher expression congruity in the mutant background (Figure 3D; Supplementary Figure 3C). Less conserved genes displayed similar patterns of de-repression in the *kmt6* mutant background but overall lower levels of transcription. This is consistent with highly conserved genes being under selection for gene expression robustness and the H3K27me3 marks acting to disrupt gene regulatory mechanisms in place.

### Feature enrichment of H3K27me3-marked genes

We investigated protein functions encoded by the H3K27m3 marked gene body. We found that H3K27me3 marked genes have shorter transcripts than unmarked genes (median 1452 *vs.* 1841 bp; *p*-value < 2.2e-16) and have a lower GC content (51% *vs.* 53%; *p*-value < 2.2e-16; Figure 4A). We found that highly conserved genes covered by H3K27me3 marks have significantly less codon usage bias than unmarked genes (CAI median of 0.778 vs. 0.785; *p*-value = 0.0002). The encoded proteins differ also in amino acid composition with an aromatic codon score median of 0.092 and 0.074 for marked and unmarked genes, respectively. We evaluated thermodynamic properties related to protein interaction and folding. Marked genes have higher amino acid hydrophobicity compared to unmarked genes (median of −0.264 vs −0.426; Figure 4A). Conserved genes with H3K27me3 marks are enriched for functions related to oxidation-reduction, transmembrane functions, and transcriptional regulation (*p*-value < 0.0001; Figure 4B, Supplementary Figure 4A). Highly conserved unmarked genes are enriched for housekeeping functions such as cellular metabolic and macromolecular processes (Supplementary Figure 4B).

**Figure 4.**
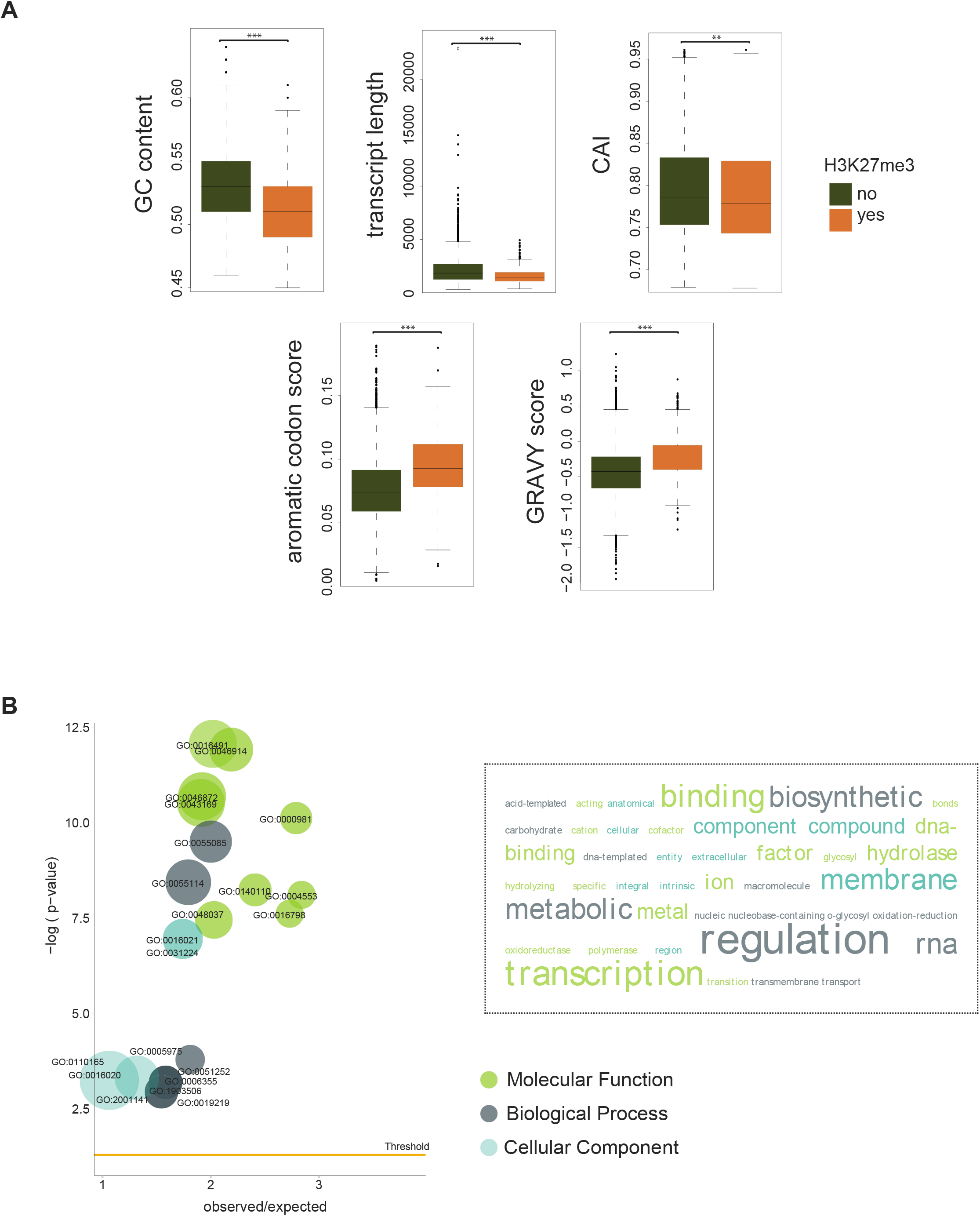
Features of H3K27me3-marked genes. A) Analyses of codon and amino acid features of highly conserved genes in *Fusarium graminearum*. Green and orange colors indicate the presence or absence of H3K27me3 in the gene body. CAI refers to the Codon Adaptation Index, GRAVY scores amino acid hydrophobicity. B) Gene ontology (GO) term enrichment analysis of highly conserved genes marked by H3K27me3. The word size represents the enriched GO term size. Threshold line refers to p-value < 0.0001.

### Evolutionary conservation of H3K27me3 marks among *Fusarium* fungi

To understand the evolutionary longevity of epigenetic effects on gene expression robustness, we analyzed H3K27me3 marks across the genomes of distantly related *Fusarium* species. We examined 3727 (*Fusarium* conserved) highly conserved single copy orthologs shared by *F. graminearum, F. oxysporum* and *F. fujikuroi.* As expected, the large majority of the highly conserved genes are not covered by histone H3K27me3 marks (*n* = 3253; Figure 5A). Genes marked by H3K27me3 majoritarily shared the mark among orthologs (59.8%; *n* = 297/473). This suggests that H3K27me3 is often retained over significant evolutionary timescales (Figure 5A). Next, we asked if H3K27me3 mark conservation in *Fusarium* is correlated with transcriptional variation. Using ranked gene expression, we compared gene transcription levels among *F. oxysporum, F. fujikuroi* and *F. graminearum*, as well as members of the FGSC during host infection and in culture condition. H3K27me3 marked genes in *Fusarium* show strong and consistent transcriptional repression across species and conditions (Figure 5B). Interestingly, genes with conserved H3K27me3 marks show also conserved patterns of up-regulation during infection. This is despite the fact that *F. graminearum* and *F. oxysporum* assays have been carried out on different hosts (wheat and tomato, respectively). Conversely, in culture conditions H3K27me3 marked genes of *F. oxysporum* and *F. fujikuroi* show stronger repression compared to *F. graminearum.* Overall, gene expression of H3K27me3 marked genes was consistently lower in culture conditions compared to host infection highlighting the de-repression caused by infection stress (Figure 5B).

**Figure 5.**
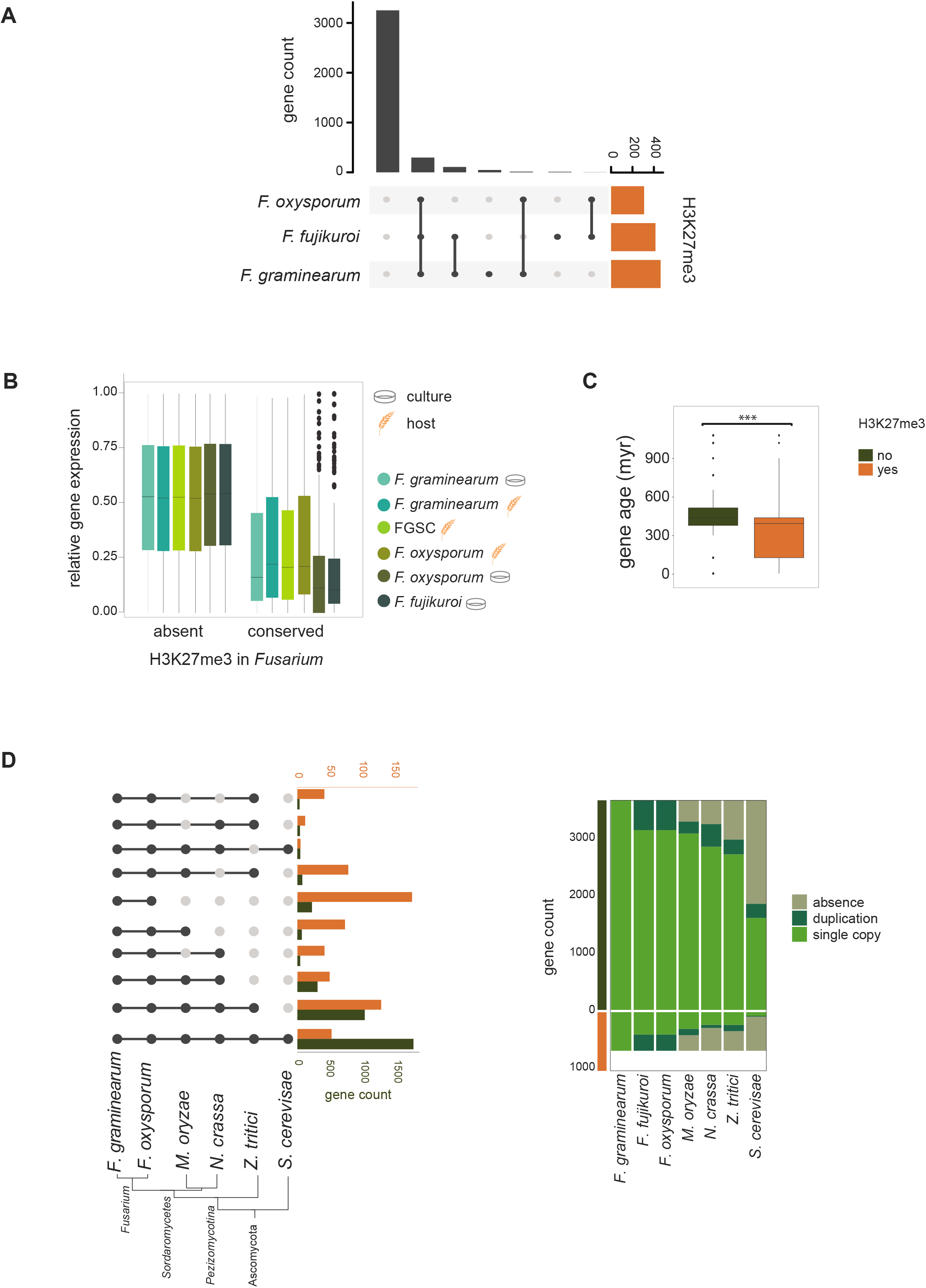
Conservation of H3K27me3 gene body occupancy in highly conserved genes across different species in the *Fusarium* genus. A) Co-occurrence of H3K27me3 marks in *F. graminearum, F. oxysporum* and *F. fujikuroi.* B) Relative gene expression (ranked) of *Fusarium* species under different growth conditions with or without H3K27me3 marks. Host and culture medium describe the two environmental conditions. C) Age inference of the highly conserved gene set. D) Highly conserved orthologues among ascomycetes. The phylogenetic tree is based on single-copy orthologues. Orange and green colors indicate the presence and absence of H3K27me3 in the gene body, respectively.

### Evolutionary origins of highly conserved genes

To investigate the evolutionary origins of highly conserved genes within FGSC (*n* = 4332), we expanded the orthology analyses to a set of representative ascomycetes. We found a striking difference in the evolutionary age of H3K27me3 marked genes compared to unmarked genes (Figure 5D). The majority of the unmarked genes are shared among ascomycetes. Conversely, H3K27me3 marked genes were largely restricted to the *Fusarium* genus. Based on gene age calibrations for *F. oxysporum*, we find that unmarked highly conserved genes have a median age of ~439 million years. H3K27me3 marked genes have a median age of ~394 million years (Figure 5C) matching the predicted age of the *Fusarium* genus (Summerell et al. 2010). Genes marked by H3K27me3 showed a much higher duplication rate compared to unmarked genes (Figure 5D). Gene duplication events can trigger genomic defense mechanisms such as repeat-induced point mutations (RIP). Hence, we analyzed evidence for RIP-like mutations in the gene sets of *F. graminearum* but found only a small portion of the genome to be affected (<1.1%). We found no difference between the extent of RIP-like mutations in H3K27me3 marked versus unmarked genes (Supplementary Figure 5).

### Impact of H3K27me3 marks on gene dispensability

Repressive histone marks can have an impact on gene dispensability. We first analyzed a dataset of 657 transcription factor deletion mutant lines screened across 17 phenotypic traits using the *F. graminearum* PH1 strain as a genetic background (Son et al 2011). Out of the 657 transcription factors, 238 are considered highly conserved within FGSC based on our classification. Highly conserved H3K27me3 marked genes encoding transcription factors show lower transcription levels, lower expression congruity and lower protein sequence conservation compared to marked genes (Supplementary Figure 6B). The screened phenotypes include sexual development, mycelial growth, secondary metabolite production, virulence, and stress response. As previously reported by Son et al. (2011), only 26% of the deletion mutants show detectable phenotypic effects. Mutants with phenotypic effects related to sexual development carried mostly (94.4%) a knock-out of an unmarked gene. Whereas altered phenotypes associated with virulence and stress response were only found in unmarked gene mutants (Figure 6). Expanding to the full data set (highly conserved + variable genes; n=657 genes), the same pattern was retained (Supplementary figure 6A). Next, we analyzed a dataset of 101 *F. oxysporum f. sp. lycopersici* mutants with reduced pathogenicity compared to the wild type (Michielse et al. 2008). Similarly, out of all genes with phenotypic effects only 7% (7/101) were marked with H3K27me3. Among the highly conserved genes, only one gene (1/36) was marked by H3K27me3 (Supplementary Figure 6B). Hence, our analysis shows that H3K27me3 marks are linked to gene dispensability in the *Fusarium* genus.

**Figure 6.**
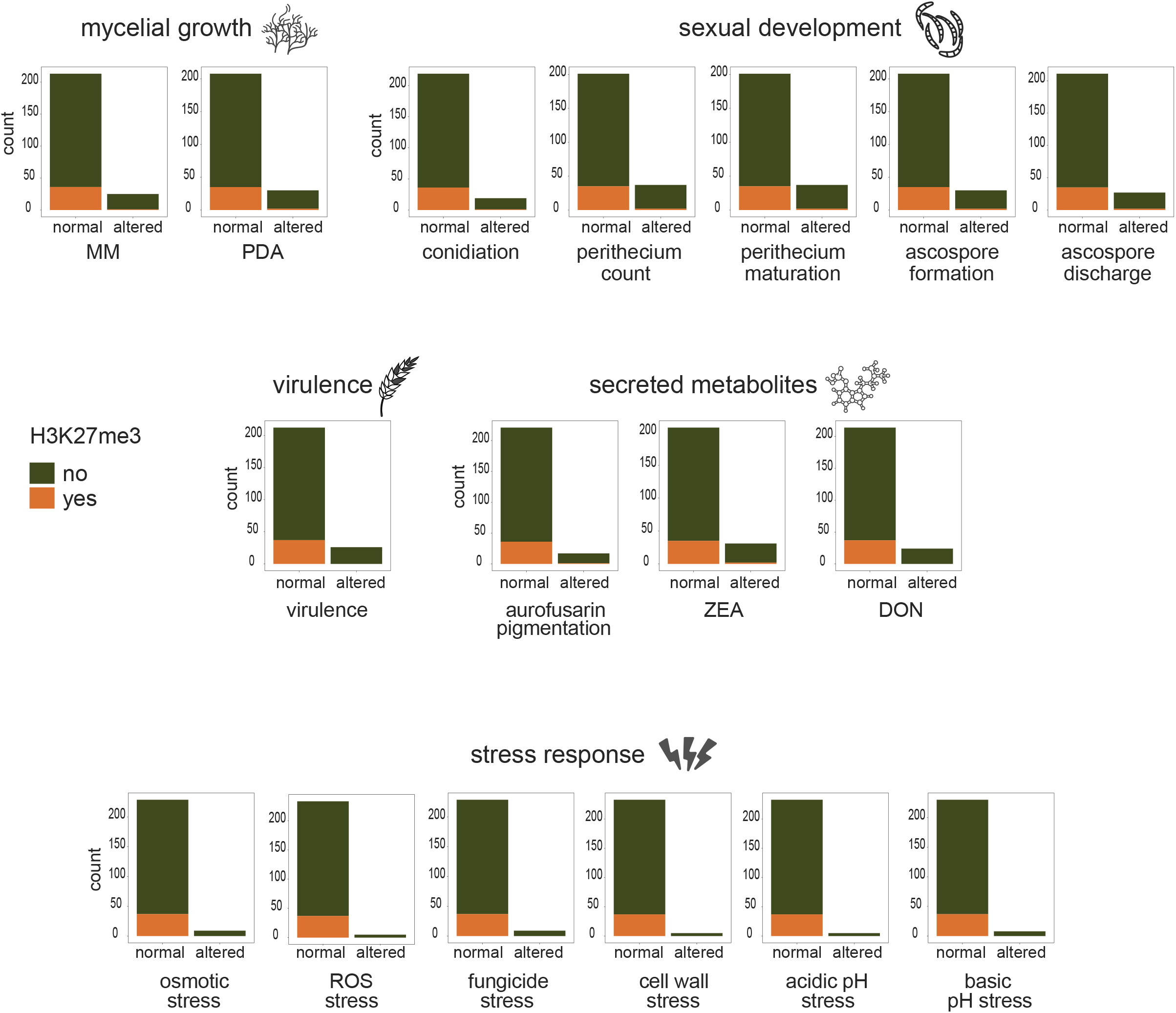
Phenotypic effect analyses of transcription factor deletion mutants in the *F. graminearum* reference isolate PH1 background. Normal refers to mutants without any overt phenotypic change compared to the wild type. Green and orange colors indicate the presence or absence of H3K27me3 in the gene body. The presented data is a subset of the dataset by Son et al. 2011 filtered for highly conserved genes. DON – deoxynivalenol, ZEA-zearalenone.

### Linking H3K27me3 marks to protein conservation in filamentous fungi

To investigate repressive histone marks and protein conservation over deep evolutionary timespans, we analyzed single-copy orthologues (*n* = 4034) shared between the distantly related model fungi *N. crassa, M. oryzae* and *Z. tritici.* Only a single ortholog had conserved H3K27me3 marks among the species (Figure 7A). Overall, we found 111 genes with H3K27me3 marks in at least one of the species (Figure 7B). The majority (55.4%) of all orthologues were also highly conserved in *Fusarium* and 38% of H3K27me3 marked genes in any of the distant fungi show also H3K27me3 marks in *F. graminearum* (Figure 7B). However, only 44/111 of the H3K27me3 marked genes were highly conserved in *Fusarium.* Hence, protein conservation is reduced in *Fusarium* for genes with repressive histone marks. We also investigated links between H3K27me3 marked genes among distant fungi and transcriptional responses in different environments (host and culture condition) among members of the FGSC (Figure 7C). We selected genes, which were both conserved among distant fungi and in *Fusarium (n* = 2213/4034). Genes with shared H3K27me3 marks among distant fungi and *Fusarium* showed lower gene expression and high expression robustness across environments (Figure 7C, Supplementary Figure 7). Interestingly, highly conserved genes with H3K27me3 marks in distant fungi but not in *Fusarium* share similar sequence conservation levels but have lower expression robustness in FGSC (Figure 7C). Together, our findings show that highly conserved genes with repressive H3K27me3 marks in distantly related fungi underpin transcriptional perturbation at microevolutionary scale in the species complex.

**Figure 7.**
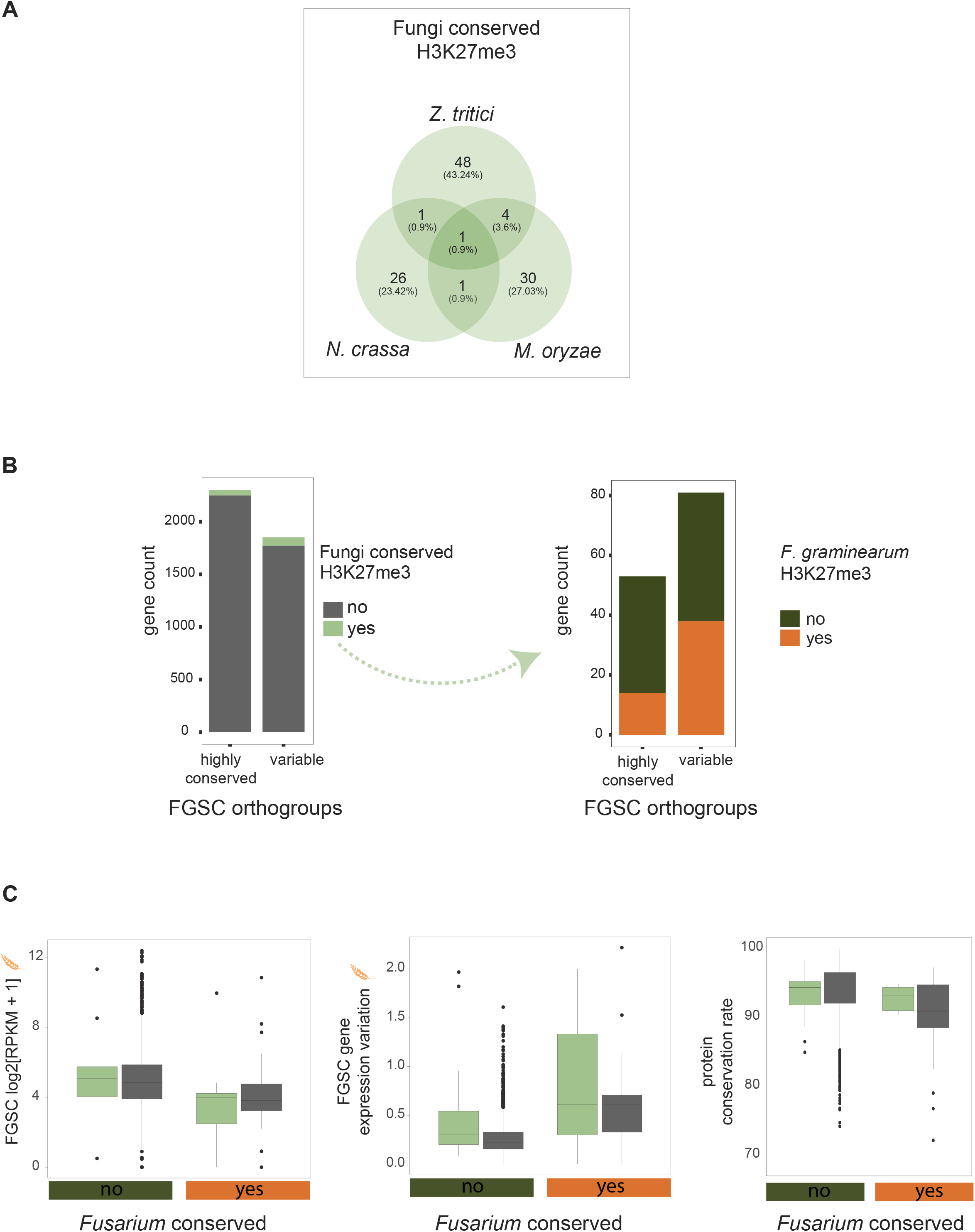
H3K27me3 histone methylation conservation among ascomycetes. A) Single-copy orthologues shared between *Neurospora crassa, Magnaporthe oryzae* and *Zymoseptoria tritici* with overlapping H3K27me3 gene body occupancy between species. B) Orthologues found in the FGSC according to *Fusarium* genus conservation categories (highly conserved vs. variable). Subsequent green arrow and bar charts refer to a subset of the data including genes marked by H3K27me3 (*n* = 111) with shared orthologues in the *Fusarium* genus. C) Gene expression analyses for host condition, expression robustness and protein sequence conservation rate of highly conserved genes in the FGSC. H3K27me3 mark conservation was assessed among distant ascomycetes.

## Discussion

We analyzed a major group of plant pathogenic fungi to understand factors associated with gene expression robustness over deeper evolutionary timescales. We performed transcriptome-wide analyses to assess expression patterns across environments and species. Integrating histone marks data across the genus and additional ascomycetes, we established predictors for expression robustness, protein conservation and gene dispensability.

### Repressive histone marks predict gene expression and dispensability

Highly conserved genes tend to be constitutively expressed to maintain basic cellular functions (She et al. 2009). We show that highly conserved genes marked by H3K27me3 deviate from these archetypical housekeeping properties. Compared to unmarked genes, marked genes exhibit shorter transcripts, lower GC content, and weaker codon usage bias, which are features associated with fast evolving genes (Lipman et al. 2002; Zhang and Yang 2015). Functions encoded by marked genes were also enriched for transcriptional regulation and binding functions. This is consistent with findings in the *Arabidopsis* genus with an enrichment for similar functions (Ha et al. 2011). The higher levels of aromatic codons and hydrophobicity of proteins encoded by marked genes suggest weakened selection against misfolding and aggregation (Bastolla et al. 2004). In contrast, highly conserved genes are typically under strong purifying selection to encode key functions in molecular and cellular processes. Similarly, highly expressed genes are under strong selection for expression robustness to reduce translational errors, protein misfolding and to avoid promiscuous protein-protein interactions (Zhang and Yang 2015; Agozzino and Dill 2018). Destabilized proteins can cause protein-protein aggregation with deleterious consequences for cellular functions including membrane integrity (Stefani and Dobson 2003). Our analyses show that highly conserved genes marked by H3K27me3 encode distinct functions compared to similarly conserved genes without heterochromatin marks. In conjunction, our analyses show that H3K27me3 marked genes are under relaxed selection pressure to avoid protein cytotoxicity.

We show that transcription factor deletion mutants affecting highly conserved genes marked by H3K27me3 are less likely to generate an overt phenotypic defect compared to unmarked but similarly conserved genes. A lack of apparent phenotypic consequences could stem from genetic redundancy. However, our analysis focused only on single copy orthologs making gene duplications an unlikely explanation (*i.e.* by creating redundancy). Alternatively, genes marked by H3K27me3 could have recently lost essentiality either through changes in the environmental interactions of the organism or mutations in the genetic background creating redundancy (Bergmiller et al. 2012). Interestingly, essential genes marked by H3K27me3 can become dispensable through epistatic interactions, monogenic suppressors or similar mechanisms (Li et al. 2019; Rousset et al. 2021). Genomic regions with repressive histone marks may coincide with higher densities of repetitive elements. In some fungi, such regions are targeted by genomic defense mechanisms including RIP (Baez-Ortega et al.) causing loss-of-function mutations (Möller et al. 2021) We found no evidence for higher rates of RIP-like mutations in marked versus unmarked genes. Hence, gene dispensability most likely evolved through other mechanisms.

### H3K27me3 perturbs gene expression robustness and congruity of highly expressed genes

Highly conserved genes in the *Fusarium* genus show strong positive associations between gene expression and degree of conservation. Hence, our results confirm the E-R correlation observed broadly among eukaryotes stipulating that purifying selection acts on translation efficiency (Zhang and Yang 2015) Beyond levels of expression, we found that expression robustness is strongly associated with the degree of protein conservation independent of the environment. The association likely reflects selection acting on regulatory elements to retain expression homeostasis under external and internal perturbations (Kitano 2004). H3K27me3 plays a significant role in gene expression robustness among closely related species. Less conserved genes covered by H3K27me3 are largely transcriptionally silent confirming well-documented patterns in fungi (Freitag 2017) and other eukaryotes (Zhang and Yang 2015). We find that highly conserved genes covered by H3K27me3 suffer mostly partial transcriptional repression and rarely complete silencing. Hence, the genes retained translation proficiency despite the repressive nature of the histone modifications. Robustness typically evolves in order to tolerate environmental perturbations and facilitates the evolvability of complex systems (Kitano 2004; Liu et al. 2020). We found that highly conserved genes with repressive histone marks showed a much higher level of gene expression variation and lower expression congruity compared to unmarked genes. Associations of histone modifications and gene expression across species was found previously in primates (Zhou et al. 2014). However, much more extensive work was done on cytosine DNA methylation. Interestingly, conservation of gene body methylation in coding sequences is positively correlated with both gene expression levels and slower protein sequence evolution, but negatively correlated with gene expression robustness (Chuang and Chiang 2014; Seymour and Gaut 2020).

We found that genes with repressive histone marks showed a weaker association between protein conservation and transcriptional robustness compared to unmarked genes. The loss in expression variation could be a consequence of polymorphism in H3K27me3 marks within the species complex. However, our phylogenetic analyses showed that gene body H3K27me3 marks of highly conserved genes are largely maintained. Interestingly, the deletion of the methyltransferase KMT6 responsible for deposition of H3K27me3 (Connolly et al. 2013) restored gene expression of marked genes to similar levels as highly conserved unmarked genes. Hence, there is an additional layer of transcriptional regulation and maintenance of translation efficiency in highly conserved genes. After depletion of H3K27me3 in the KMT6 mutant, highly conserved genes marked in the wild type showed a strong increase in gene expression congruity similar to unmarked genes. Hence, H3K27me3 seems to be the causal factor disturbing otherwise similar expression congruity of highly conserved genes. Our analyses also indicate that the sensitivity to transcriptional perturbation by H3K27me3 marks within the species complex is likely shared among distantly related ascomycetes.

Our study establishes evolutionary links between expression robustness and protein evolution mediated by repressive histone modifications. Highly conserved genes marked by H3K27me3 are transcriptionally active despite the repressive nature of the mark but suffer from perturbed expression robustness compared to unmarked genes. Highly conserved marked genes show enrichment in environmental stress related functions, carry hallmarks of fast evolving genes and, hence, do not follow the housekeeping gene archetype. We show that H3K27me3 marks can blur the general E-R correlation of sequence conservation and expression levels. Hence, histone modifications are a key link between protein evolvability and gene essentiality.

## Acknowledgements

We thank Dirk Balmer at Syngenta Inc. and the CBS-KNAW culture collection for providing fungal material. Ursula Oggenfus and Vinciane Mossion provided helpful comments on the manuscript. The research was supported by FAPESP (Fundação de Amparo à Pesquisa do Estado de São Paulo) grant process 2019/16045-0. DC received support from the Swiss National Science Foundation (grant 31003A_173265).

