## Supplementary Figures for "Histone H3K27 methylation perturbs transcriptional robustness and underpins dispensability of highly conserved genes in fungi"

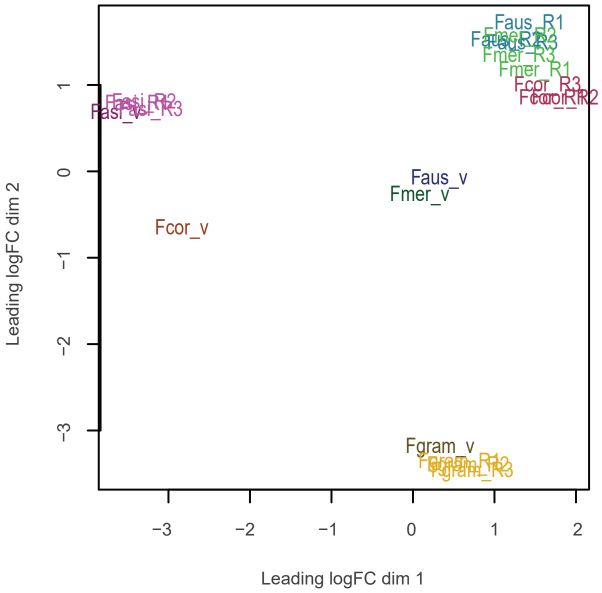


**Supplementary Figure S1**: Multi-dimensional scaling analysis of the transcriptomic dataset of the *Fusarium graminearum* species complex. Samples with suffix _v refer to *in vitro* condition single replicate, _R to host infection, followed by the replicate number. Fgram refers to *F. graminearum*, Fcor to *F. cortaderiae*, Faus to *F. austroamericanum*, Fmer to *F. meridionale* and Fasi to *F. asiaticum*.

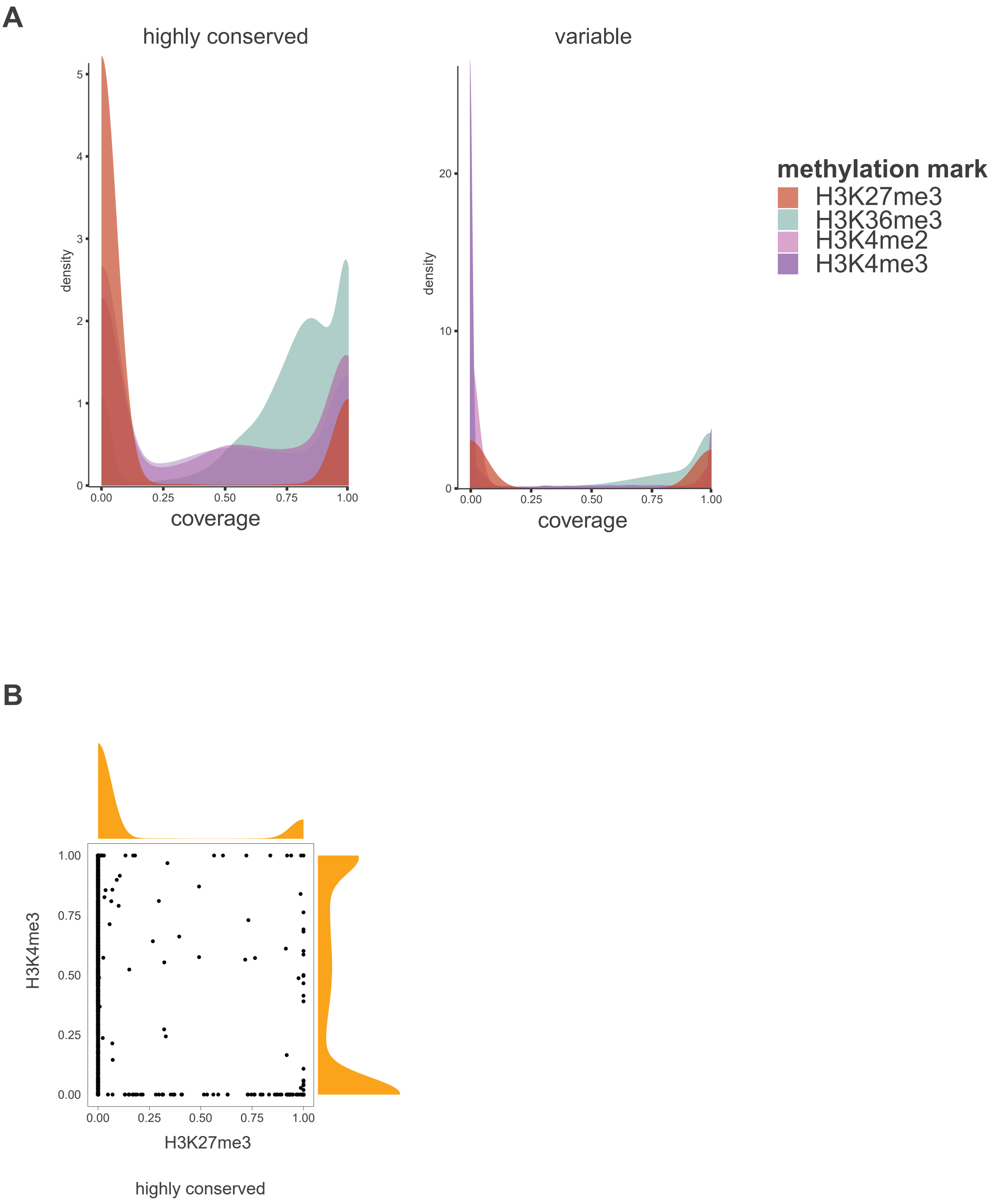


**Supplementary Figure S2:** A) Density plot of histone modification mark coverage highly conserved genes and variable genes in *F. graminearum* (PH1) (Connolly et al. 2013). B) Distribution of H3K4me3 and H3K27me3 histone marks among highly conserved genes.

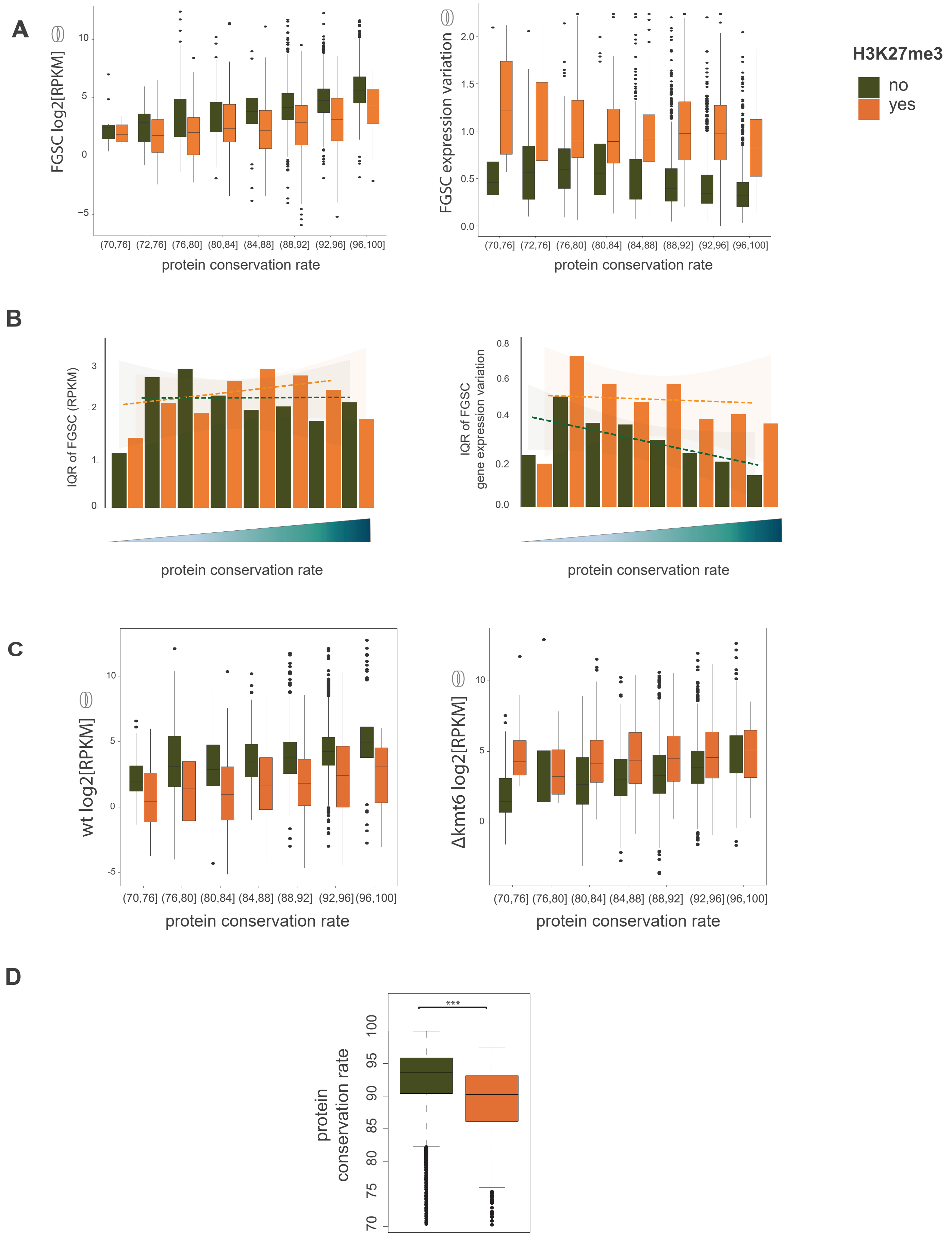


**Supplementary Figure S3:** A) Distribution of mean expression and mean expression variation (*i.e.* robustness) of highly conserved genes during *in vitro* growth based on protein sequence conservation rate. C) Gene expression congruity and gene expression variation congruity of highly conserved genes based on the interquartile range (IQR) of gene expression and gene expression variation during host infection (see also Figure 3B-C). D) Gene expression analysis of the *Fusarium graminearum* (PH1) wild type and mutant strains of highly conserved genes during *in vitro* growth. E) Distribution of protein conservation rate of highly conserved genes. Orange refers to genes marked by H3K27me3 and green to unmarked genes.

**A**

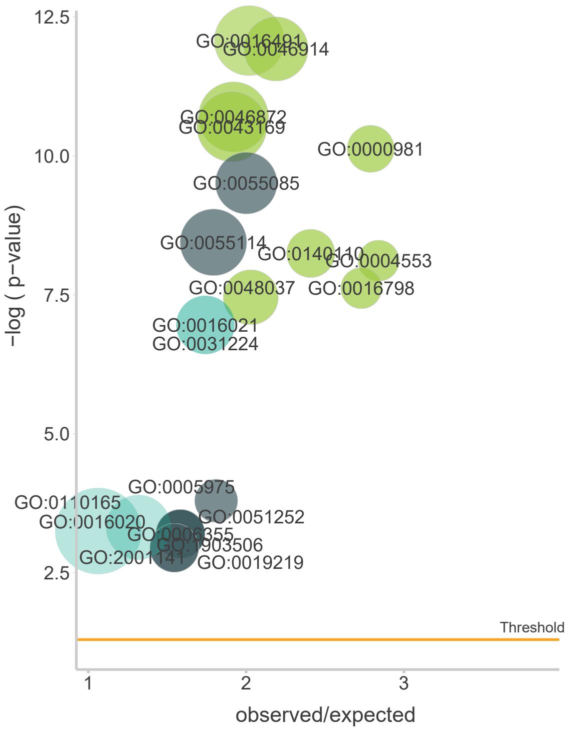


| ID | Description |
|---|---|
| GO:0016491 | oxidoreductase activity |
| GO:0046914 | transition metal ion binding |
| GO:0046872 | metal ion binding |
| GO:0043169 | cation binding |
| GO:0000981 | DNA−binding transcription factor activity |
| GO:0140110 | transcription regulator activity |
| GO:0004553 | hydrolase activity |
| GO:0016798 | hydrolase activity |
| GO:0048037 | cofactor binding |
| GO:0055085 | transmembrane transport |
| GO:0055114 | oxidation−reduction process |
| GO:0005975 | carbohydrate metabolic process |
| GO:0006355 | regulation of transcription |
| GO:0019219 | regulation of nucleobase |
| GO:0051252 | regulation of RNA metabolic process |
| GO:1903506 | regulation of nucleic acid |
| GO:2001141 | regulation of RNA biosynthetic process |
| GO:0016021 | integral component of membrane |
| GO:0031224 | intrinsic component of membrane |
| GO:0016020 | membrane |
| GO:0110165 | cellular anatomical entity |

**B**

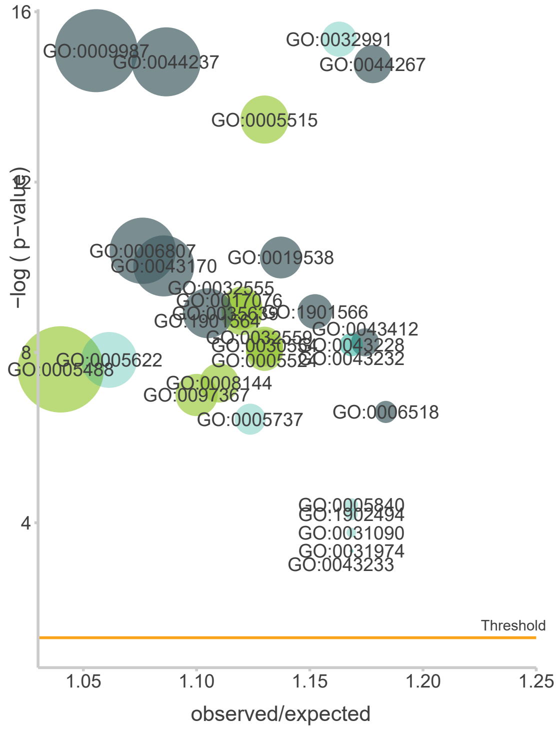


| ID | Description |
|---|---|
| GO:0005515 | protein binding |
| GO:0017076 | purine nucleotide binding |
| GO:0032555 | purine ribonucleotide binding |
| GO:0035639 | purine ribonucleoside triphosphate bindi... |
| GO:0030554 | adenyl nucleotide binding |
| GO:0032559 | adenyl ribonucleotide binding |
| GO:0005524 | ATP binding |
| GO:0005488 | binding |
| GO:0008144 | drug binding |
| GO:0097367 | carbohydrate derivative binding |
| GO:0009987 | cellular process |
| GO:0044237 | cellular metabolic process |
| GO:0044267 | cellular protein metabolic process |
| GO:0006807 | nitrogen compound metabolic process |
| GO:0019538 | protein metabolic process |
| GO:0043170 | macromolecule metabolic process |
| GO:1901566 | organonitrogen compound biosynthetic pro... |
| GO:1901564 | organonitrogen compound metabolic proces... |
| GO:0043412 | macromolecule modification |
| GO:0006518 | peptide metabolic process |
| GO:0032991 | protein−containing complex |
| GO:0043228 | non−membrane−bounded organelle |
| GO:0043232 | intracellular non−membrane−bounded organ... |
| GO:0005622 | intracellular |
| GO:0005737 | cytoplasm |
| GO:0005840 | ribosome |
| GO:1902494 | catalytic complex |
| GO:0031090 | organelle membrane |
| GO:0031974 | membrane−enclosed lumen |
| GO:0043233 | organelle lumen |

● Molecular Function ● Biological Process ● Cellular Component

**Supplementary Figure S4.** Enrichment of gene ontology terms in highly conserved genes marked (A) and unmarked (B) by H3K27me3.

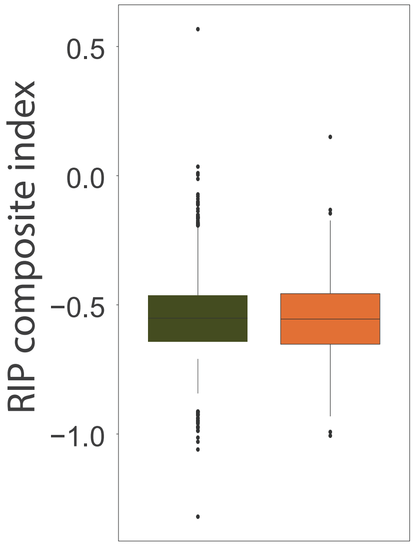


**Supplementary Figure S5:** Repeat induced point mutations (RIP) composite index in highly conserved genes marked and unmarked by H3K27me3.

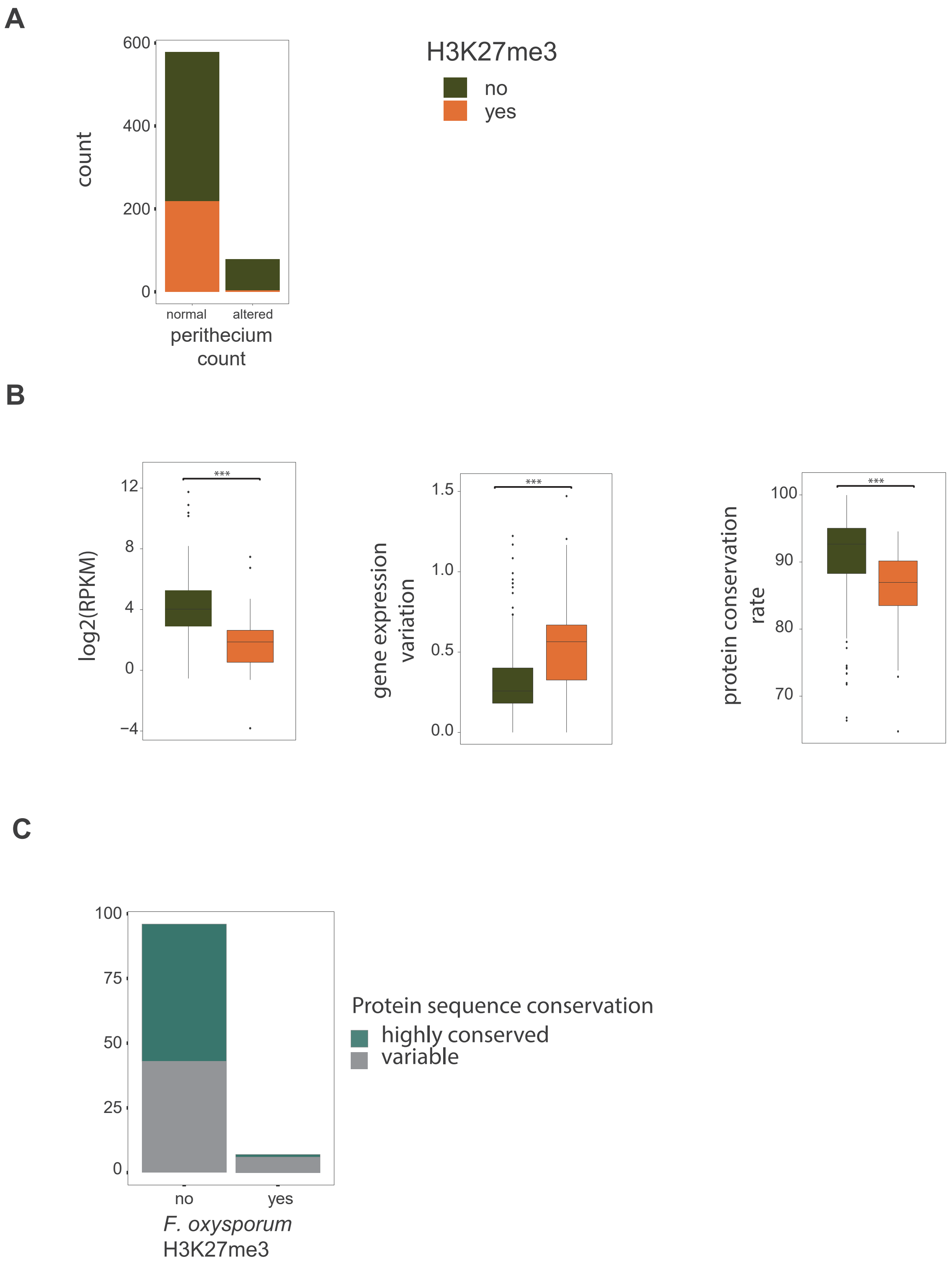


**Supplementary Figure S6.** A) Phenotypic analyses of transcription factor deletion mutants (*F. graminearum* PH1 background) assessed for perithecium counts ($n$ = 657 genes). "Normal" and "altered" phenotypes related to the wild type strain PH1. B) Gene expression, expression variation and protein conservation of highly conserved genes encoding transcription factors screened using deletion mutants (see also Figure 7A). Orange refers to genes marked by H3K27me3 and green to unmarked genes. C) *F. oxysporum f. sp. lycopersici* deletion mutants screened for loss or reduction of pathogenicity. Genes are categorized into marked and unmarked by H3K27me3.

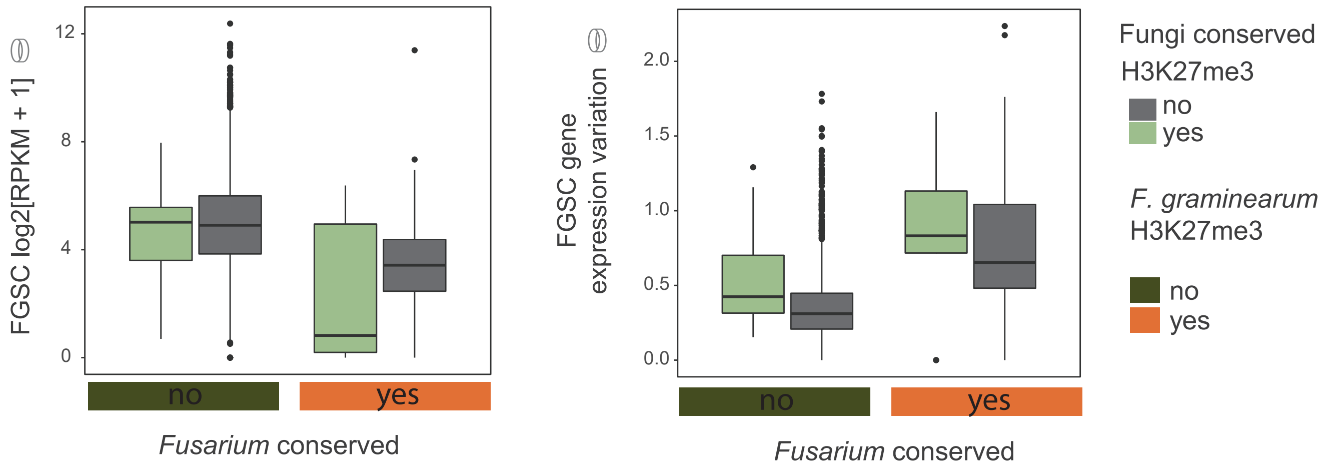


**Supplementary Figure S7.** Gene expression analyses and expression robustness for *in vitro* condition of highly conserved genes in the FGSC. H3K27me3 mark conservation was assessed among distant ascomycetes (Fungi conserved H3K27me3).
